# deepNGS Navigator: Exploring antibody NGS datasets using deep contrastive learning

**DOI:** 10.1101/2025.01.27.634805

**Authors:** Homa MohammadiPeyhani, Edith Lee, Richard Bonneau, Vladimir Gligorijevic, Jae Hyeon Lee

## Abstract

High-throughput sequencing uncovers how B-cells adapt and evolve in response to antigens by generating B-cell receptor (BCR) sequences at an unprecedented scale. As BCR datasets grow to be millions of sequences, using efficient computational methods becomes crucial for analyzing and understanding complex patterns within the data. One important aspect of antibody sequence analysis is detecting clonal families or clusters of related sequences, whether they come from immunization, synthetic libraries or even ML-generated datasets. Such analysis helps us to understand how sequences are evolutionarily connected, and how they might have been selected or evolved. Here we introduce deepNGS Navigator, a computational tool that leverages language models and contrastive learning to transform large datasets of antibody sequences into 2D representations. The resulting 2D maps offer an intuitive visualization of overall diversity of input datasets, which can be clustered based on the sequence distances and their densities across the map. Beyond grouping related sequences, the 2D maps can also point to evolutionary trajectories and capture mutational patterns among closely related sequences. By analyzing properties like charge, hydrophobicity, number of sequence neighbors, and read counts, the maps highlight which clusters are most promising for further investigation while also detecting anomalies or noisy sequences with higher risk. We demonstrate deepNGS Navigator’s utilities on various datasets, including: 1) a synthetic library from a yeast display targeting HER2, 2) a machine learning-generated dataset with a hierarchical tree structure, 3) NGS sequences from a llama immunized against COVID RBD, 4) human naive and memory B-cell sequences, and 5) an insilico dataset simulating B-cell clonal lineages.

## INTRODUCTION

Advances in high-throughput sequencing technology have significantly enhanced our ability to investigate large biological and synthetic antibody libraries, providing opportunities to harness deep insights into immune response and underlying patterns. However, analyzing and interpreting the large volume of complex data generated from immunization or synthetic B-cell libraries poses a significant challenge (1). Though phylogenetic tree based methods provide valuable insights into the clonal evolution of B-cells, their application to large-scale B-cell repertoires is limited by computational scalability, incomplete germline databases of camelids (2), complex gene conversion mechanisms in rabbits (3), and a requirement on the minimum number of sequences within each clonal group for meaningful analysis, which often restricts analyses to only the largest V(D)J gene groups (4; 5; 6; 7). Machine learning (ML) methods for two-dimensional visualization of sequences like UMAP(8) are sometimes used to enable visual exploration of sequence dataset structure (2). However, it is often unclear how to interpret and use such projection maps for practical applications, such as clonotyping and hit selection or for follow-up experimental characterizations. To address these challenges, we developed deepNGS Navigator that leverages deep learning based language models (LM) and contrastive learning to transform high-dimensional antibody sequence repertoire data into intuitive two-dimensional maps (Figure 1). This approach captures and visualizes the edit distance neighborhood structure of a dataset and provides an intuitive understanding of the dataset’s diversity. Furthermore, clustering based on these maps could potentially group sequences with related functions, facilitating follow-up studies that benefit from the selection of sequences from diverse and distinct clusters.

**Fig. 1.**
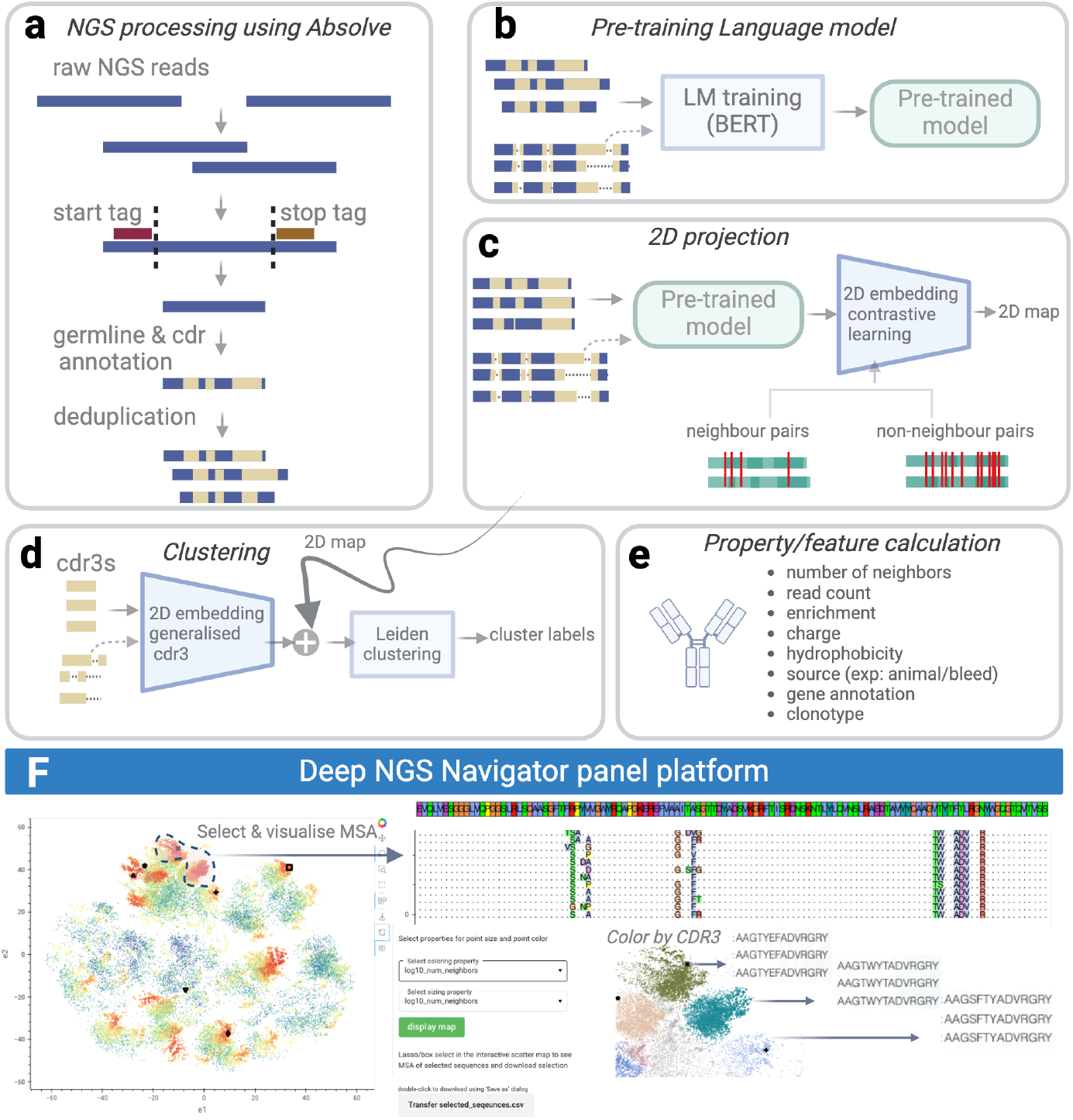
deepNGS Navigator workflow. **(a)** Preprocessing raw NGS datasets using Absolve, including merging forward and reverse reads and optionally translating them into proteins (or users can directly cluster nucleotide sequences). The sequences are annotated by their CDR3, deduplicated, and their copy number is recorded for each unique sequence. In the final table each row represents a unique sequence with minimum heavy and/or light chain and their respective CDR3 information. **(b)** Pretraining a language model on preprocessed NGS datasets by using either aligned sequences or groups of sequences with same sequence and CDR3 lengths. **(c)** 2D projecting using LM and contrastive learning, where neighbors and non-neighbors for each sequence are defined based on allowed edit distance in CDR3 and full sequence. **(d)** Assigning cluster labels using Leiden algorithm. Optionally edge cases of each cluster can be refined using CDR3 embedding of sequences. **(e)** Further annotating dataset by calculating various properties and features, including charge, hydrophobicity, and neighbor count. **(f)** Interactive visualization of final 2D map in deepNGS panel platform, allowing users to color the map based on various properties, select sequences of interest and plot their multiple sequence alignments (MSA). Created with BioRender.com

deepNGS Navigator takes NGS datasets (nucleotides or amino acids) as input (Figure 1a). It utilizes a BERT-type language model (9) to embed sequences into high dimensional feature space, which can represent complex relationships and patterns among input sequences (Figure 1b). Then using a contrastive learning technique, inspired by frameworks like SimCLR (10) and t-SimCNE (11), the high dimensional LM embedding is projected to 2D maps (Figure 1c). As part of contrastive learning process, neighbors and non-neighbors for each sequence are defined based on allowed edit distance in both the complementary-determining regions 3 (CDR3) and the full sequence, tailored specifically for antibody sequences. Next, Leiden algorithm (12) is used on the 2D projection to identify clusters. Given the importance and diversity of the CDR3s, users have an option to incorporate additional embeddings of CDR3s at a lower weight to further refine the maps (Figure 1d). Finally, users can annotate the dataset with various metrics, such as biophysical properties like charge and hydrophobicity, and then interactively explore the 2D maps within the deepNGS Panel platform. This includes drawing, selecting, visualizing MSAs, and downloading selected sequences (Fig 1. e,f). To showcase the method’s broad applicability, we used deepNGS Navigator to analyze following datasets from various studies:

1. We generated 2D maps of synthetic library sequences from a yeast display annotated by FACS binding labels to HER2. The results of the analysis show that our approach produces clusters with smaller average pairwise edit distances and lower label entropy compared to alternative methods explored.
2. We analyzed synthetic machine learning-generated sequences with a hierarchical tree structure rooted on a “seed” sequence. Our analysis shows that the deepNGS Navigator method provides an embedding map whose global organization captures the edit distance relationship better than that of alternatives.
3. We clustered NGS sequences from a llama immunized against COVID RBD and show that the method achieves fewer, larger clusters with comparable enrichment purity compared to a prior method.
4. We applied the visualization to a dataset consisting of naive and memory derived human B-cells and show the proximity of sequences in the embedding map is correlated with their potential biological relations.
5. We reconstructed clonal families in an *in silico* designed dataset and demonstrated that deepNGS performs similar to the state-of-the-art methods in accurately identifying lineages. Moreover, it shows superior performance in separating true lineages from noisy datapoints, while relying solely on sequence information without gene annotation guidance.

These applications highlight the potential of deepNGS Navigator to transform complex sequence spaces into intuitive 2D maps, that facilitate discovery workflows by clustering sequences with similar biological functions.

## MATERIALS AND METHODS

### Data Collection and Initial Processing

Pre-processing antibody NGS datasets includes the following steps: (i) merging raw forward and reverse reads, (ii) translating sequences to protein, and (iii) performing germline annotation. In this study, raw datasets were pre-processed using Absolve (13). Output tables from Absolve were further refined by correcting sequence prefixes and suffixes to reduce primer contamination based on their germline information. Then, sequences were deduplicated, while keeping read count as an attribute for number of occurences of each sequence in dataset. Germline annotation (step (iii)) can be useful for sanity checks of the final maps, but it is not mandatory. Translating sequences to proteins in step (ii) is also optional. As demonstrated in our last case study, deepNGS can effectively analyze nucleotide sequences as well.

deepNGS Navigator expects input data structured in a tabular format with each row representing a unique sequence, containing the following columns:

- fv heavy: heavy chain sequence.
- fv light: light chain sequence.
- HCDR3: heavy chain CDR3.
- LCDR3: light chain CDR3.

If one chain is missing, its respective column can be left empty. Based on the case study and available data, users can choose to apply deepNGS to all input sequences with varying lengths, resulting in a single comprehensive map for all of them. In this scenario, it is necessary to align the sequences using methods like AbAlign (14) or AHo numbering (15). Alternatively, for more granular embedding maps, users can group sequences by their sequence and CDR3 lengths, and optionally by V-gene, and generate 2D maps specific to each group.

### Language Model Training and Embedding Sequences into 2D Space

Language models (LM) have been shown to capture complex patterns in biological sequence data (16). In deepNGS Navigator, by default, a BERT type LM (9) is pre-trained on all input antibody repertoire data. When working with a small dataset, one can replace this step by using LMs pre-trained on larger, more general datasets such as AbLang (17), which is specifically trained on antibodies. Next, we use contrastive learning on high dimensional embeddings of sequences to project them to 2D maps. Specifically, we draw inspiration from SimCLR (10), and adapt the recently introduced t-SimCNE method (11), which has shown promising applications in analyzing and clustering image data.

The architecture of deepNGS Navigator model consists of three main parts:

- A LM component with BERT-style architecture, initialized from a pre-trained model, which generates high dimensional embeddings of each sequence (with a default size of 256). As a result, regardless of the original sequence length, the component maps all sequences to a fixed-size vector.
- A linear transformation layer that accepts the LM embeddings as input and outputs embeddings of the same dimension.
- A final linear projection head that projects the high dimensional embeddings to a 2-dimensional space.

### Neighbor Definition

Our contrastive learning methods require the definition of positive and negative sequence pairs. Here, neighbors and non-neighbors. In this work, we define neighbors based on Hamming distance. Sequences within a defined Hamming distance cutoff are considered neighbors. Users can either set aHamming distance cutoff or suggest an estimated number of neighbors that each sequence should have on average. This estimate is then used to calculate the appropriate Hamming distance cutoff. This is especially useful when analyzing datasets with high diversity where it is challenging to define reasonable cutoffs in advance. For an efficient clustering using contrastive learning regardless of the dataset, we recommend each sequence to have an average of at least 5 to 10 neighbors. Achieving this number of neighbors may require a different Hamming distance cutoff depending on the dataset. We suggest starting with this range and adjusting the cutoff as needed, either tightening or relaxing it based on the results. In antibodies, CDR3 similarity is particularly important for maintaining functional relevance. To take this into account and control the CDR3 diversity among neighbors, we set a separate threshold on Hamming distance specifically for the CDR3 region. This threshold can be defined manually or adapted on a dataset basis as mentioned above. The default Hamming distance cutoff is 5 for the full sequence, with a maximum 2 out of 5 in the CDR3 region, unless specified by user input. As an important note, sequences that do not meet the minimum requirement of having at least 5 neighbors are excluded from training by default. This exclusion criterion is helpful as sequences outside of it have difficult-to-identify placements on the map and often tend to obscure relations of other sequences with more neighbors by acting as background noise.

### Optimizing Loss Function for 2D Embeddings

We utilized the Cauchy similarity loss inspired by t-SimCNE with a few modifications that allow users to experiment and adjust depending on the case study. Given a batch of *N* sequences 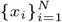, we embed each sequence into a *d*-dimensional latent embedding *e*_*i*_ *∈* ℝ ^*d*^ using a LM *f* : *x*_*i*_ *→ e*_*i*_. For each sequence *x*_*i*_, we randomly select and embed a single neighbor 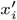 from its precomputed list of neighbors, resulting in 2*N* embedded sequences.

Next, we compute the following pairwise Euclidean distances between embeddings:

1. 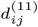: the squared Euclidean distances between all pairs of original sequences, defined as:

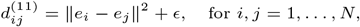

where *ϵ* is a small constant added for numerical stability.
2. 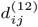: the squared Euclidean distances between each sequence *e*_*i*_ and selected neighbor sequences 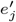, given by:

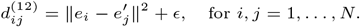

These two distance matrices are then concatenated to form a combined distance matrix *d*_sq_ *∈* ℝ^*N ×*2*N*^ :

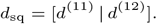

We then convert the distances into Cauchy similarities *ϕ*_*ij*_, where:

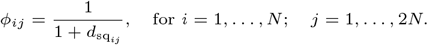

Optionally, the distance matrix can be modified using a fitted kernel transformation (18), as: 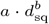 where *a* and *b* are defined by the following curve-fitting process: First, depending on *min dist* parameter (with a default value of 1.0) a target function *f* (*x, min_dist*) assigns a value of 1 to distances below *min dist* threshold and decays exponentially for larger values. The model function 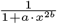 is then fitted to this target function using curve fit module from scipy library (19), optimizing *a* and *b*. Once fitted, the transformation 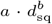 is applied to the distance matrix in Cauchy similarities *ϕ*_*ij*_ equation above. This adjustment preserves connections for small distances and controls the decay for larger ones, enhancing how relationships are represented in projection maps.

To correctly apply repulsive forces only between non-neighbors, we construct a mask matrix *M ∈ {*0, 1*}*^*N ×*2*N*^ :

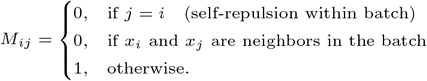

Finally, the t-SimCNE loss consists of two main components:

Positive Loss.

The positive term encourages sequences to be close to their corresponding neighbors:

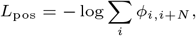

Negative Loss.

The negative term ensures that sequences are repelled from all other non-neighboring sequences:

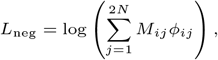

where the mask *M*_*ij*_ ensures that repulsion is only applied between non-neighboring sequences.

Thus, the overall loss function is a sum of the positive and negative terms:

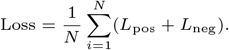

In addition, users can experiment with the option to normalize the loss by number of neighbors and non-neighbors. This option normalizes positive loss by the number of neighbors and negative loss by the number of non-neighbors for each sequence within a batch. This normalization helps in extreme cases where a sequence has few or no neighbors. It ensures the model does not overly focus on positioning that particular sequence either exceptionally close or far from the rest, maintaining a balanced perspective.

### Data loader and Training

We used the PyTorch WeightedRandomSampler as our sampling algorithm, which samples from the dataset based on predefined probabilities of each data point. We computed sampling weight as the logarithm of the number of neighbors. This approach prioritizes sequences with higher number of neighbors during sampling and led to improved model performance compared to uniform random sampling.

### Training Procedure

The training process is structured in three stages as suggested by t-SimCNE authors. In the first stage, both the LM and the linear transformation layer are trained together to better adapt the pre-trained model to input sequences (default 400 epochs). In the second stage, the LM and linear transformation are frozen, and only the projection head is trained (default 100 epochs). Finally, in the third stage, the entire model is unfrozen and trained further (default 400 epochs).

### Clustering and Refinement

To identify clusters on the final map, Leiden algorithm is applied to 2D embeddings. To further refine boundaries and edge cases, for instance, when sequences on the embedding map have an equal probability of belonging to multiple clusters, we prioritize CDR3 similarity to decide on their cluster labels. This is implemented using DenseClus method (20), where two separate embeddings of the same dataset are combined by taking their sum to form their union. In our case, the primary 2D embedding is derived from contrastive learning and a secondary embedding represents the features of the CDR3 of sequences. To create the CDR3 features, we first reduce their diversity by categorizing amino acids based on properties like charge and hydrophobicity (generalized CDR3). We then break the generalized CDR3s into k-mer fragments, reduce their dimensionality with PCA, and apply UMAP to achieve a standard 2D representation.

### Visualization and Interactive Exploration

The resulting data is visualized through a web browser based Panel (21) platform, allowing users to interactively explore the 2D maps generated by deepNGS Navigator. Currently, the platform supports visualizing the final projection map, allowing users to customize the coloring scheme and point size based on various dataset properties, such as CDR annotations, number of neighbors, sequence length, and gene annotations. In addition, users can explore and zoom into different parts of the map, and use lasso or box select tools to choose sequences of interest. These selected sequences can then be visualized using the platform’s integrated MSA viewer or downloaded directly in a table format for further analysis.

## RESULTS

### Clustering antibody sequences from yeast display libraries with FACS labels

Clustering NGS datasets in a way that reflects their relationships and functionality is an important step for efficient hit discovery. If clusters correctly represent the functionality of sequences, we can prioritize diverse selections across map in the initial rounds and shift the focus on optimizing best binders by mining the most promising clusters in subsequent stages. We benchmarked the performance of deepNGS Navigator using 100,000 sampled sequences from Minot et al.’s yeast display libraries targeting HER2 (22). This dataset, labeled by fluorescence-activated cell sorting (FACS) and deep sequencing, provides an ideal dataset for such analysis. In addition to deepNGS Navigator, which trains a custom language model on the input dataset, we explored the impact of contrastive learning on clustering by using the same pre-trained language model with dimensionality reduction techniques like t-SNE (23) and UMAP. In addition, we evaluated a generalized pre-trained model, AbLang (17), combined with UMAP to compare with deepNGS pre-trained LM on input data. As shown in Figure 2b, while all methods showed some degree of sensitivity to binding properties of sequences, deepNGS Navigator and LM+t-SNE methods more accurately distinguish between binders, weak binders, and non-binders. To quantify these embedding qualities, we used Leiden algorithm (12) for clustering them, applying 20 neighbors as the hyperparameter across all methods. The resulting number of clusters were: 100 clusters for deepNGS, 125 in LM+t-SNE, and 50 clusters for both LM +UMAP and AbLang+UMAP (Figure 2c). We evaluated edit distance of sequences and entropy of FACS labels within each cluster, where their overall distributions per method is shown in Figure 2d,e. Ideally, clusters should exhibit minimized edit distance and entropy while globally separating FACS-derived labels across clusters. deepNGS Navigator consistently demonstrated this trend, validating its high-resolution and fine clustering performance both within and across clusters. This benchmark illustrates how different clustering methods can lead to different interpretation of the same sequences space. deepNGS Navigator generated an intuitive map that separates sequences based on their binding functionality while minimizing edit distances within clusters, such map facilitates achieving the primary goal of identifying diverse binders from large datasets. This approach is adaptable to other datasets, where sequences whether single, paired chains, or specific regions like H3 and L3, can be embedded and clustered, producing maps that reflect their biological relatedness and functionality. This enables users to prioritize diverse sequence selections across promising clusters based on different factors. For example some binder enriched clusters are at the same time highly charged, posing a risk of non-specificity. Such factors can be systematically profiled across entire map, guiding strategic sequence selection and experiments.

**Fig. 2.**
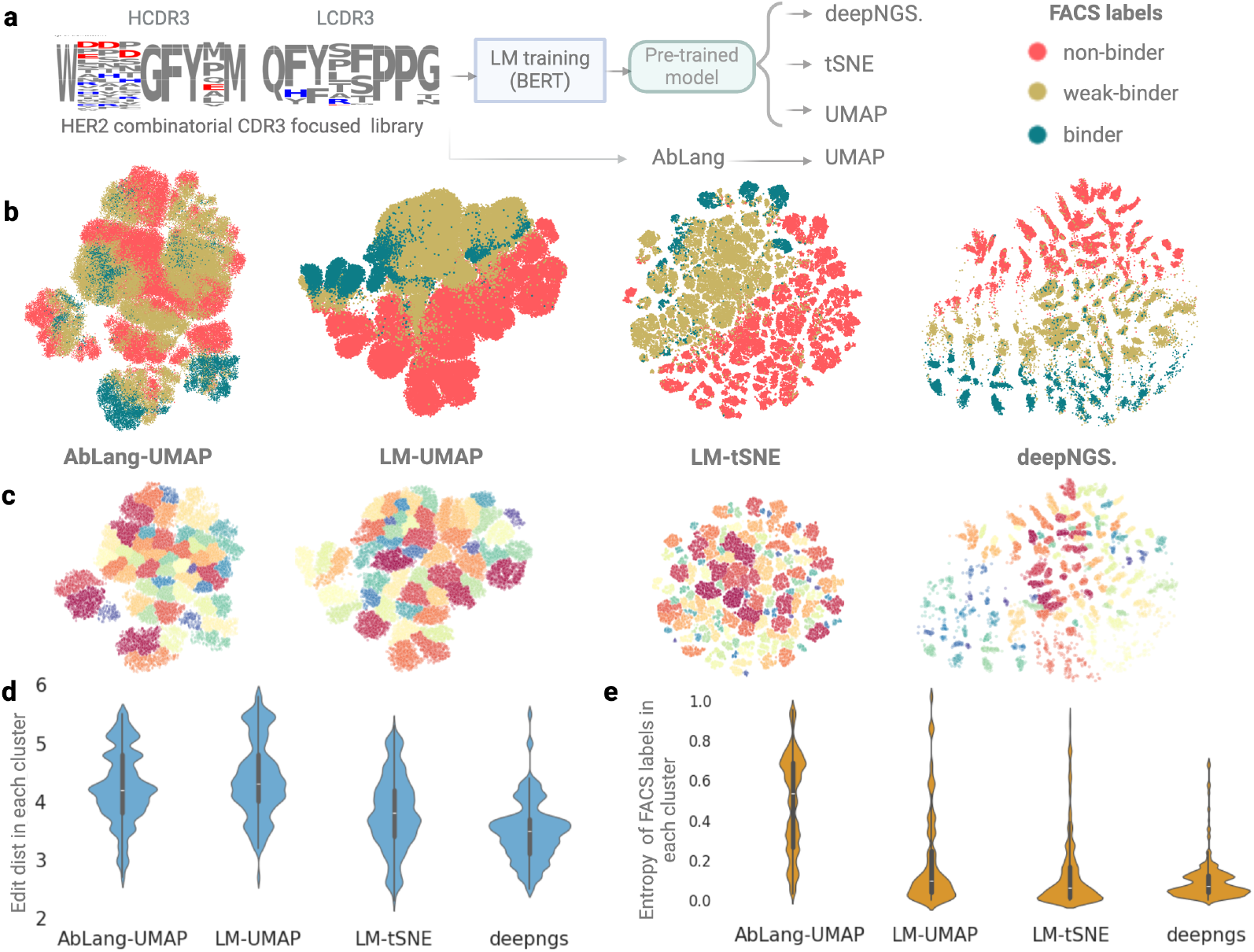
**(a)** Comparison of clustering methods on the Her2 yeast display dataset. **(b)** Visualizes the binding FACS labels on 2D projections. **(c)** Displays the grouped sequences using the Leiden algorithm. The quality of clusters is assessed based on intra-cluster. **(d)** edit distance and **(e)** FACS labels entropy. Created with BioRender.com

### Evaluating clustering performance on a synthetic dataset with hierarchical structure

To investigate how well the global structure of a dataset is mirrored in a two-dimensional space, we applied deepNGS Navigator to a synthetically designed heavy chain dataset with a known generation logic. This dataset from Lin Li et al (24) originates from a single seed sequence and expands to approximately 60,000 variants through random mutation and advanced ML generative methods including Gaussian Process and Ensemble models, with hill climb, genetic and Gibbs sampling algorithms, denoted as GP-HC, GP-GA, GP-Gibbs, En-HC, En-GA and En-Gibbs(24). This dataset’s hierarchical structure is reminiscent of an evolutionary tree, with the seed sequence at the root, the sequences with few small random mutations forming immediate branches, and then the designed sequences representing more distant branches. This structure allows us to assess whether our clustering algorithm can effectively distinguish sequences of varying relatedness, akin to differentiating the root and branches of a tree. Beside deepNGS Navigator, we evaluated the performance of alternative techniques as described in the previous section, including deepNGS Navigator’s pre-trained LM combined with t-SNE or UMAP, and AbLang combined with UMAP. In Figure 3, we compared final maps both qualitatively by coloring sequences with their origin, and quantitatively by analyzing the correlation between their edit distances and Euclidean distances. As shown in Figure 3a-d, while all methods provide sequence embedding maps of similar global organization, they differed in resolution and accuracy. First, the customized language model in deepNGS Navigator shows more sensitivity to separation of sequences with different origins compared to general antibody LM in AbLang. Furthermore, with a correlation score of 0.94 between the edit distance from the seed and the Euclidean distance on the map from the seed, deepNGS Navigator 2d embedding offers a finer ordering based on a gradual increase in number of mutations. It organizes the seed sequence at the top, positions the random mutant variants around it, and progressively maps sequences designed using the Gaussian Process (GP) method—which have lower edit distances to the seed—extending outward. The most distant sequences were the ensemble (En) designs, reflecting greater divergence. This spatial arrangement aligns with the expected hierarchical relationships among the sequences. Projecting high dimensional space of antibody sequences into 2D maps, while balancing both global and local structure, is crucial for producing a meaningful low dimensional representation of it, however challenging. While it is important to capture broad biological patterns at a global level, the local neighbourhoods and trajectories of sequences should reflect fine details, such as specific sequence motifs, and gradual increase of mutational load. In this case study, we demonstrated that deepNGS Navigator effectively maintains both global and local structures within the dataset by clearly separating sequences based on their origin and showing a strong correlation between the edit distance from the seed sequence and the Euclidean distance on the map. This allows meaningful exploration and exploitation both inside and across different clusters.

**Fig. 3.**
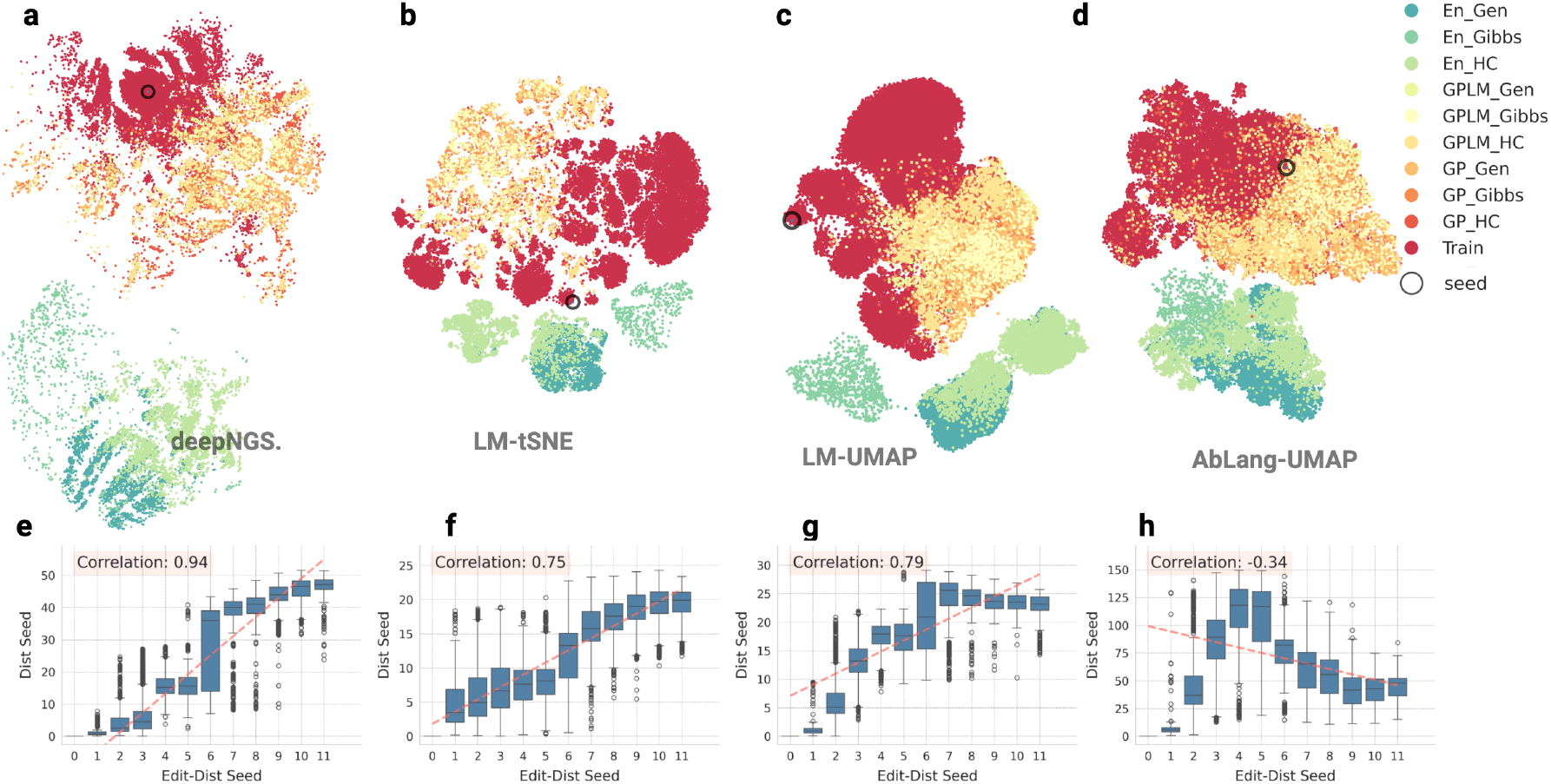
Comparison of different clustering tools on representing a hierarchical synthetic dataset. **(a-d)** show the 2d embedding colored by origin of sequences, starting with the seed sequence, followed by random low-mutation variants (Train), and designs generated by Gaussian Process (GP) or Ensemble (En) methods, combined with sampling strategies such as hill climbing (HC), genetic algorithm (GA), and Gibbs. **(e-h)** shows for each method, the correlation between the Euclidean distance from the seed on the map and the edit distance from the seed (ranging from 0 to 11) for each sequence. Created with BioRender.com

### Mining antibody phage display pools generated by panning

Display technologies are a crucial element in the hit discovery pipeline for selecting highly specific antibodies. Through multiple panning rounds and conditions sequences that bind to a specific target are enriched. High throughput sequencing (HTS) allows characterizing both final and intermediate panning round sequences. To interpret and leverage NGS with biopanning in depth, efficient computational methods are needed. In this regard, we benchmarked deepNGS Navigator and SeqUMAP using a panning dataset from Leo Hanke et al. (2) This dataset includes approximately 96,000 unique antibody sequences obtained from llama immunization targeting the SARS-CoV-2 receptor-binding domain (RBD). Our goal was to identify which tool offers a more accurate and efficient approach for identifying functionally relevant antibody sequences. This is particularly important given the limitations posed by incomplete llama germline databases, which restrict the application of methods like phylogenic tree construction or clonotyping (2). As a result, ML based methods that can infer relatedness directly from the data are valuable. SeqUMAP uses a k-mer-based approach to transform sequences into a format suitable for UMAP visualization; here we used seqUMAP with a k-mer size of 3 to analyse dataset. To accommodate variable sequence lengths as input for deepNGS, the sequences were first aligned using AbAlign (14). Both SeqUMAP and deepNGS Navigator then generated 2D visual maps of the antibody repertoire, with the sequences color-coded according to RBD enrichment levels, where enrichment per sequence is defined as the log ratio of post-to pre-panning frequency, regularized with a pseudocount (Figure 4a). To assess how effectively each method separated enriched and depleted sequences, we first classified each sequence as enriched (*enrichment >* 0.05), depleted (*enrichment <* 0.05), or unclear based on their enrichment values, then we applied Leiden clustering algorithm (12) with 10 neighbors to the 2D embeddings of both methods. SeqUMAP generated approximately 1,800 clusters, with the largest containing around 300 sequences, while deepNGS Navigator produced 505 clusters, with the largest including more than 1,300 sequences. For each cluster, we calculated the homogeneity as the ratio of the dominant enrichment label to the total cluster size. Among the top 10 largest clusters, deepNGS Navigator identified 8 predominantly enriched and 2 depleted clusters. In contrast, seqUMAP’s top 10 clusters included only 1 enriched and 6 depleted clusters, with the remaining clusters displaying mixed enrichment-depletion states. Additionally, a scatter plot showing correlation of cluster size and homogeneity indicates that deepNGS Navigator maintained higher purity despite fewer total number of clusters. In general, larger, fewer clusters are preferred as long as the resulting clusters retain high functional homogeneity. This example highlights deepNGS Navigator’s potential to achieve greater balance in this regard compared to prior methods.

**Fig. 4.**
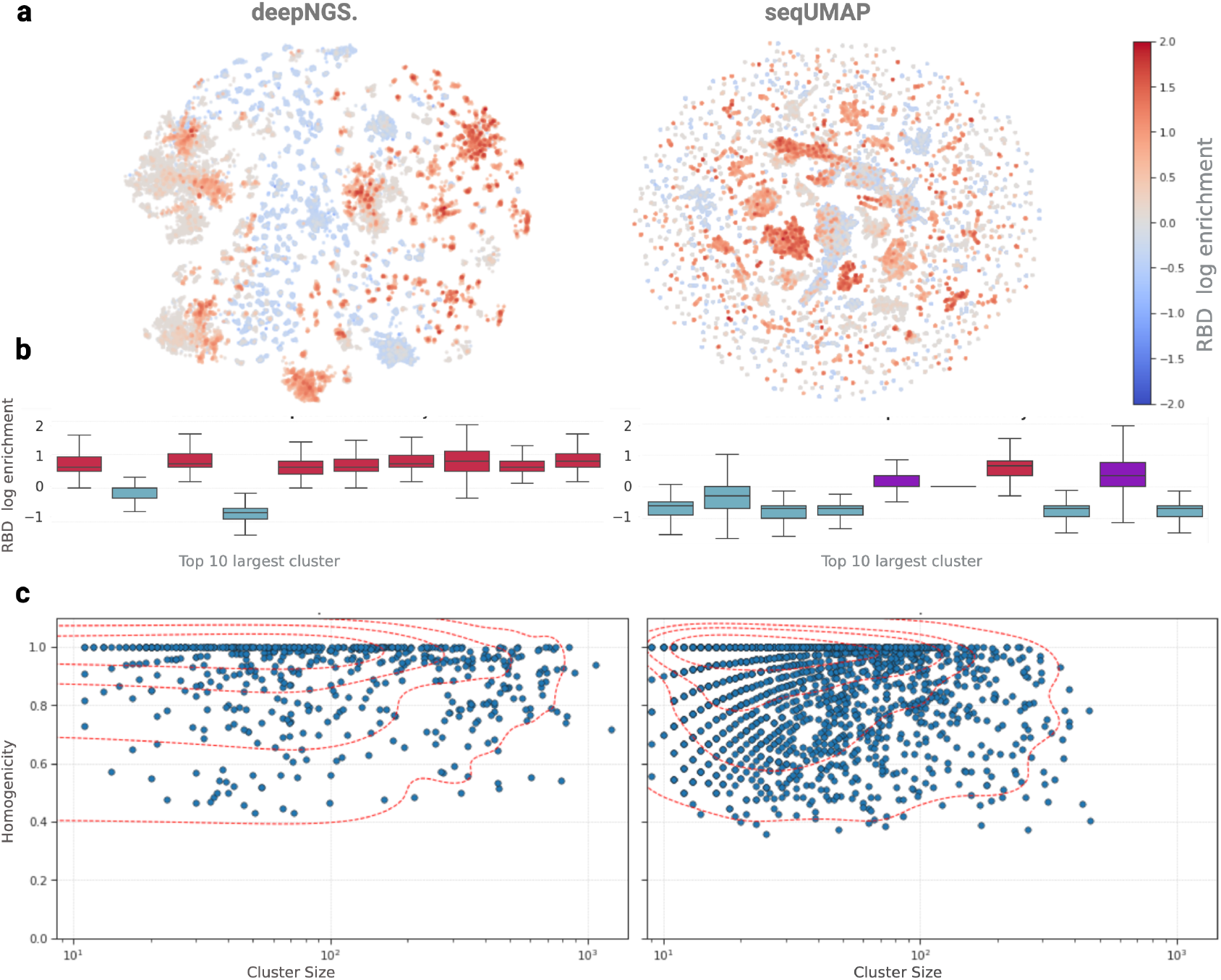
Repertoire analysis using deepNGS Navigator and seqUMAP. **(a)** 2D embedding maps generated by each method. **(b)** Enrichment of depletion profile of 10 largest clusters per method. **(c)** Correlation between cluster size and its homogeneity in terms of depletion or enrichment. Created with BioRender.com

### Differentiating Naive and Memory B Cell Subpopulations

Naive B-cells possess a broad range of antigen specificity with moderate affinity. They serve as a reservoir for generating diverse antibody responses upon initial exposures to pathogens and their Fv regions are typically representing germline diversity with the original V(D)J gene sequences without significant mutations. On the other side, Memory B-cells are generated from activated B-cells that have responded to an antigen during a primary immune response, and as a result their Fv regions have undergone somatic hypermutation (SHM), introducing point mutations in the variable regions. To investigate how effectively the characteristics of these two B-cell types are captured by deepNGS Navigator, we collected 150,000 sequences from the study by Ghraichy et al (25), representing the largest group of sequences with the same length (98 amino acids, starting from HCDR1). As shown in Figure 5a,b, deepNGS Navigator map captures separation and gradual expansion of naive to memory B-cells, highlighting the gradual increase in mutations underlying evolutionary transition of B-cell types. Furthermore inside clusters, sequences are sorted based on their V gene and J gene (Figure 5c,e), with those having higher number of neighbors in the center, expanding outward to sequences with fewer neighbors (Figure 5d). This potentially indicates sequences are ordered based on their evolutionary trajectories.

**Fig. 5.**
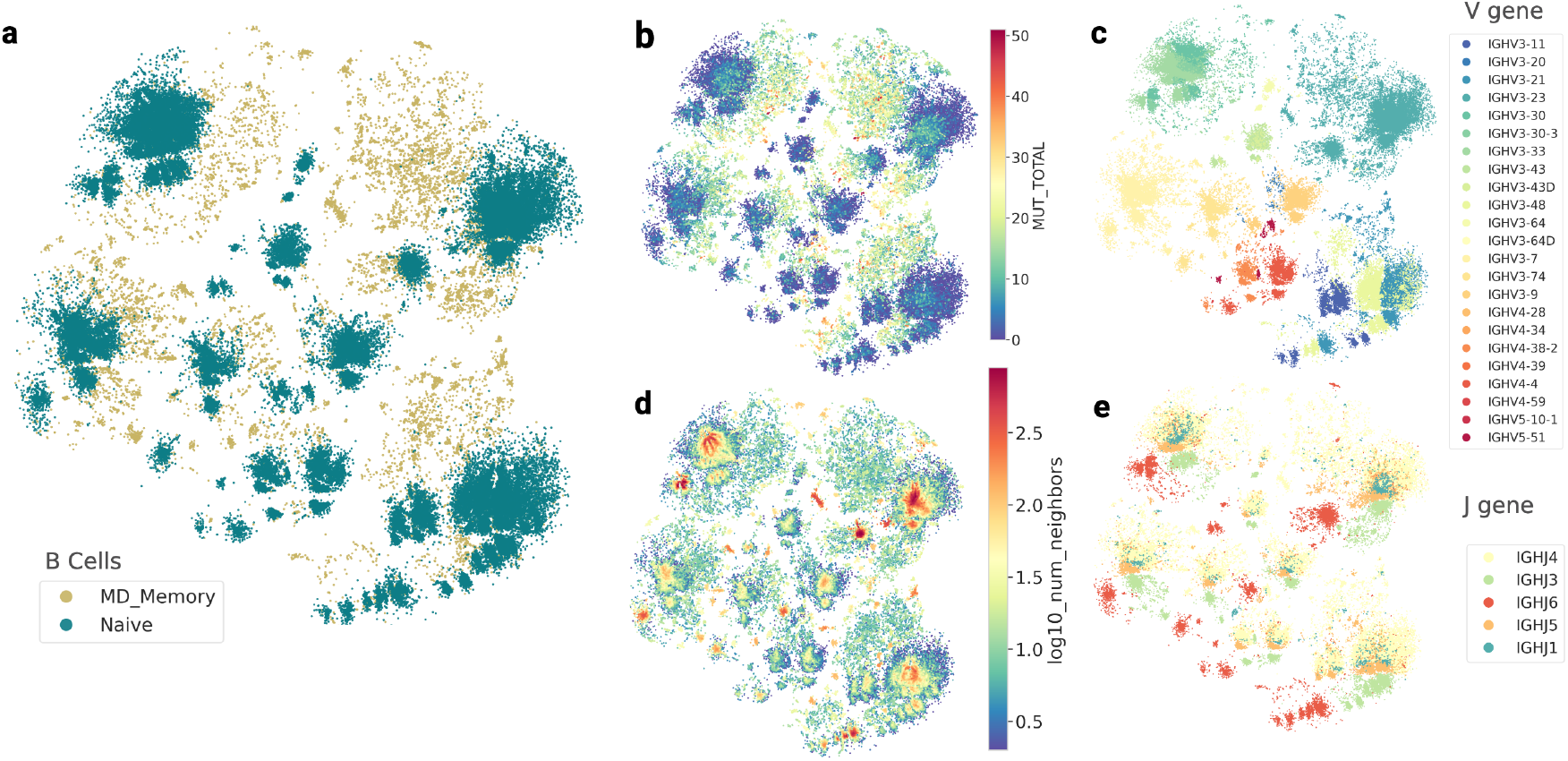
2D projection of deepNGS Navigator map annotated by: **(a)** B-cell type, **(b)** number of mutations, **(c)** V gene, **(d)** number of neighbors and, **(e)** J gene distribution. Created with BioRender.com

Using deepNGS Navigator we show the generated dimensionality-reduced projection of the B-cell repertoire preserves the key characteristics of B-cell sequences, allowing to distinguish between naïve and memory B-cell subpopulations. It has been discussed in prior studies that incorporating V and J gene usage, CDR3 physicochemical properties and global repertoire features can similarly distinguish between naïve and memory B-cell subpopulations (25). Notably, in this case, the input was limited to raw sequences, without any information about genes or additional feature guidance. This suggests, model derived meaningful insights from sequences themselves to generate informative embedding map. In addition to B-cell subpopulation discrimination, the sequence trajectories correlated with their biological properties. For instance, there is a clear separation of sequences with the same V gene family (IGHV3, IGHV4, and IGHV5) and specific genes, which are further sorted by J gene usage. This demonstrates the ability of deepNGS Navigator to capture biologically relevant patterns solely from sequence data.

### Clonal family inference from large-scale simulated B-cell lineages

The reconstruction of clonal families is a crucial step for understanding adaptive immune response and guiding drug design. During affinity maturation, B-cells originating from a unique V(D)J gene arrangement are expanded and diversified through somatic hypermutation (SHM), each variant competing for stronger binding to an antigen. However, accurately identifying these clonal families in highly diverse B-cell repertoires is a significant challenge. Current methods struggle to scale with large datasets and in addition depend on correct V(D)J gene annotation; however many germline databases are incomplete, limiting their application (2). In a study by Wang et al.,(26) a set of in silico datasets were designed to benchmark reconstruction of B cell clonal families using the state of the art methods including: FastBCR (26), MobiLLe (27), SCOPe (4), Partis (28), and VJCDR326 (29). The simulated datasets includes sequences from different size of lineages generated by various mutation rates, and to make the dataset more realistic, thousands of noise sequences (singletons) were added to each lineage. These datasets resemble clonal expansion and provides a basis for evaluating how well computational methods can identify clonal families (26). We used largest and most complex set from this study, containing about 40,000 sequences: 20,000 noise sequences and 20,000 forming 100 lineages generated at the highest mutation rate (0.01). This dataset was used to benchmark the performance of deepNGS Navigator in reconstructing clonal families, comparing it to other methods (Figure 6.a).

**Fig. 6.**
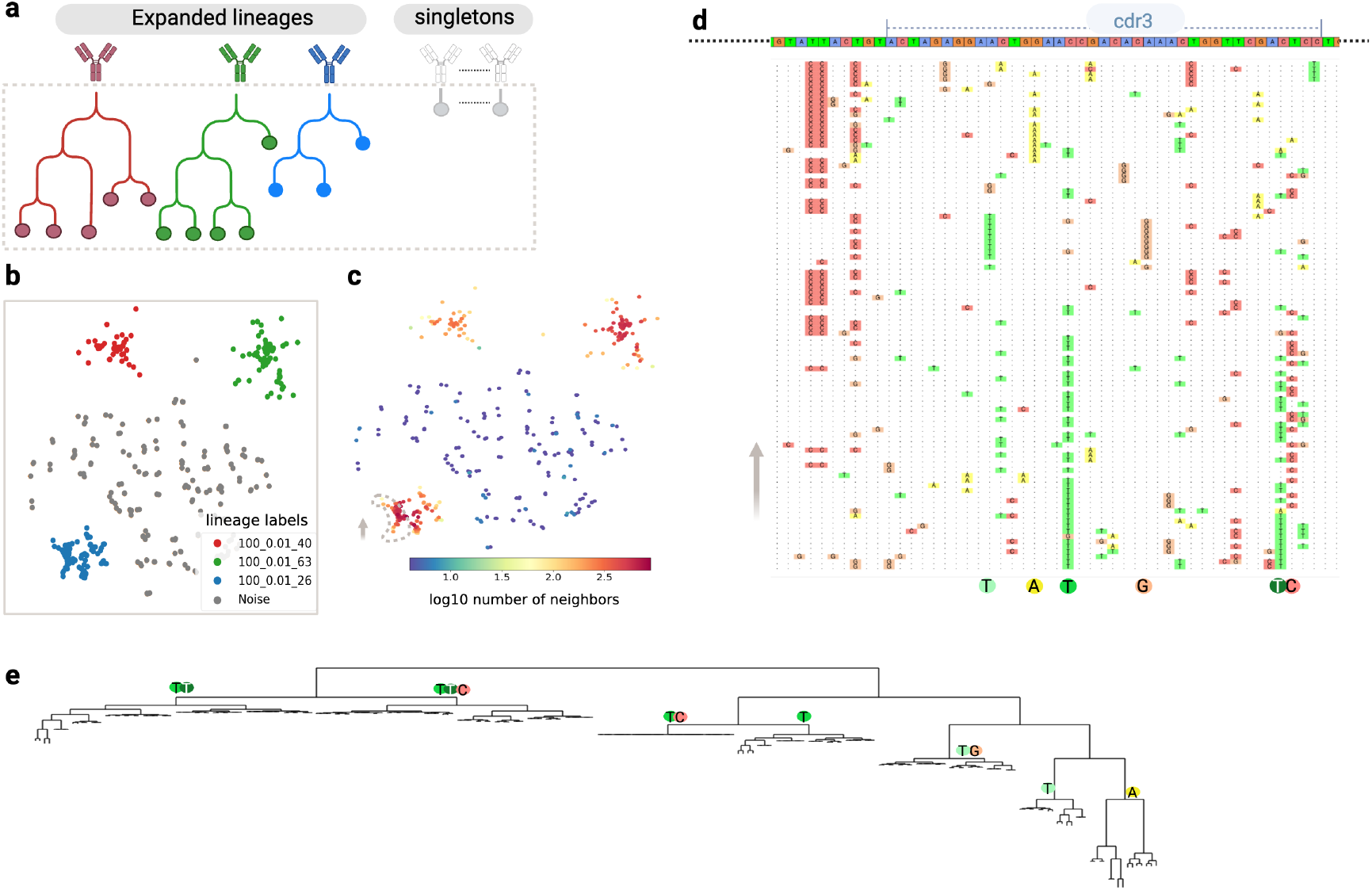
Reconstruction of clonal families from an in silico-generated dataset. **(a)** Schematic representation of the dataset, consisting of 100 lineages with added singleton sequences as noise. Sequences are grouped by total length and CDR3 length before being analyzed using deepNGS Navigator. **(b**,**c)** Example output showing sequences of length 367 and CDR3 length of 45, with a 2D embedding map colored by the original family lineage and number of neighbors each sequence has(logarithm base 10). **(d)** A view of lineage 100-0.01-26, illustrating MSA of the end of FW2 and CDR3 regions for 100 sample sequences along the vertical trajectory marked by a dashed line. The first row displays the consensus sequence, and in subsequent rows deviations from the consensus for each sequence are colored. Key conserved mutations are indicated below the MSA, showing critical mutations that distinguish sub-clusters within the lineage. **(d)** Construction of a kd-tree using the coordinates of deepNGS map for lineage 100-0.01-26, with CDR3 key mutations mapped onto it. Created with BioRender.com

Unlike previous case studies in this work, where we used protein sequences as input, this dataset consists of nucleotides, allowing us to explore deepNGS navigator’s potential in analysing DNA sequences. We grouped sequences based on their total length and CDR3 length and applied deepNGS Navigator to each group (31 length groups in total), resulting to 2d maps per length group that further clustered using Leiden method integrated in deepNGS. For example, in Figure 6, part (b) shows the embedding map for all sequences with a length of 361 and a CDR3 length of 39, color coded by their original family labels, and part (c) shows the same map color coded based on the number of neighbors (logarithm base 10) each sequence has. Using integrated Leiden algorithm we can detect three main clusters with a high number of neighbors, corresponding to the three actual families present in the group. Additionally, some sparse, isolated sequences with few neighbors which match noise sequences. Across all length groups, We compared computed cluster labels with the real labels, and calculated performance measures as described in Wang et al.’s paper (26). To assess the clustering quality of sequences with the same label (excluding noise), we used pairwise metrics across all length groups and identified clusters. A pair was defined as true positive (TP) if both sequences were correctly clustered together in both inferred and original clusters. Similarly, true negatives (TN), false positives (FP), and false negatives (FN) were defined to calculate precision, recall and F1. deepNGS Navigator achieved a high performance, with a precision of 0.99, recall of 0.94, and F1 score of 0.96, matching the best state-of-the-art results (precision:0.99, recall:0.93 and F1:0.97 respectively) without using gene annotation for guidance and instead relying only on sequences.

Furthermore, to specifically assess how well the noise data is handled, clusters categorised into three types: true-inferred clusters (TIC), mix clusters, and noise clusters. True-inferred clusters were those that reconstructed at least 80% of the true family members. Noise clusters consisted solely of noise sequences, while mix clusters were any remaining clusters(26).

With a TIC fraction of 0.98, a noise cluster fraction of 0.01, and 0.01 mixed clusters, deepNGS Navigator outperforms state-of-the-art methods (TIC: 0.88, mix:0.12 and noise:0.00); by producing pure clusters of related sequences while isolating noise sequences. We further focused on a specific lineage (100-0.01-26) as an example, and generated multiple sequence alignment (MSA) along the trajectory of the embedding map (Figure 6.d). In this lineage, diversity was concentrated towards the end of FW2 and in the CDR3 region, as visualized in Figure 6.d. The MSA reveals that as we move along the trajectory, certain patterns remain conserved, indicating that not only different lineages form distinct clusters within the map, but also that meaningful mutational patterns can be identified inside one lineage. Using the embedding coordinates, we constructed a k-d tree (30) (2) to visualize a phylogeny-like structure of sequences in this lineage (Figure 6.e). By analyzing the consensus at level 3 of this tree, we confirmed that in each sub cluster along the trajectories distinct key residues were consistently preserved. This example demonstrates that embedding maps can be directly utilized to identify closely related sequences or to construct a phylogenetic tree to visualize sequence relationships.

## DISCUSSION

B-cells are key players of adaptive immune system. They can bind and respond to a wide range of antigens by diversifying their receptors through V(D)J gene recombination and further somatic hypermutation (SHM), creating clonal families of B-cells with varying affinity to the antigen (7). Identifying these clonal families is a crucial step in drug discovery (7). Advances in high throughput sequencing (HTS) by generating B-cell receptor (BCR) sequences at an unprecedented scale has provided a much broader perspective on clonal diversity, allowing deeper analysis of immune responses. However, this massive data calls for efficient computational tools capable of unlocking their complexity. To systematically analyze these large scale sequence datasets, we leveraged deep learning based language models (BERT type) that have been shown to be exceptionally good in capturing complex patterns and extracting meaningful insights from large datasets. Then to distill the information from the language model into intuitive 2D map, we applied contrastive learning technique inspired by SimCLR (10) and t-SimCNE (11) as a method of dimensionality reduction which we customised for antibodies. The resulting maps provide an intuitive way to visualize and understand the diversity and structure of the dataset, helping to detect mutational patterns and prioritize promising candidates. The pipeline, from training the language models to contrastive learning, generating maps and interactive visualizations with various features, is all integrated into an open-source tool deepNGS Navigator and can be easily accessed by researchers, clinicians, and industry partners.

deepNGS Navigator adapts the contrastive learning by defining positive and negative pairs tailored to the unique characteristics of antibodies, such that users can control allowed diversity in CDR3 regions separately from full sequence. This customization allows for improved precision around the critical CDR3 regions, capturing subtle but yet important differences in these regions that might be overlooked by algorithms not designed specifically for antibodies.

A better understanding of clonal diversity and sequence relationships in B-cell repertoires can significantly accelerate the hit discovery process by drawing attention to clusters that show stronger antigen responses (31). These clusters can be detected in various ways depending on the experimental setting. For example, in a bio-panning experiment, clusters with higher enrichment scores will stand out (32). Similarly, in FACS experiments (33), clusters that contain mostly binders can be prioritized. While in an immunization campaign (32), embedding naive repertoires alongside sequences from immunized can highlight key differences. Or in case of multiple animal immunizations, embedding sequences from different animals together can help detect any convergence evolution among them. We benchmarked deepNGS Navigator’s performance across several case studies. In one example, using sequences from phage panning of a llama immunized against COVID RBD (2), we showed deepNGS Navigator’s clusters retain consistent enrichment or depletion patterns. It outperformed prior methods by producing fewer, larger, and more coherent clusters. In another case study, we applied deepNGS Navigator to embed and cluster sequences of a synthetic library from a yeast display annotated with FACS binding labels to HER2 (22). The resulting map efficiently separated sequences based on their binding labels while maintaining lower pairwise edit distances and reduced binding label entropy compared to alternative methods. Once promising clusters are identified, a diverse selection representing these clusters will be made to fully leverage the potential of the entire dataset. At the same time, this approach can help to filter out noisy or risky sequences that deviate significantly from the rest of the repertoire, reducing the chance of selecting unreliable candidates.

Understanding the close relatives of sequences within each cluster can also further promote selecting candidates with fewer liabilities from the beginning of hit discovery. Since, focusing only on binding affinity may overlook potential issues, such as charge patches or hydrophobicity, which could complicate optimization in later stages (24). These properties may conflict with improving affinity, leading to more challenging adjustments. By considering these factors early on, we can save time and resources in the optimization process (34). We suggest expanding and integrating other metrics into deepNGS Navigator maps such as immunogenicity risk to optimise hit selection across wider range of properties that are important for the project.

deepNGS Navigator’s application is not limited to experimentally derived sequences. With the rise of advanced generative machine learning algorithms (ML), the focus is shifting toward designing *in silico* sequences conditioned on binding to specific antigens and optimizing other biophysical properties. These ML methods can generate vast numbers of sequence designs, raising the question of how to select a diverse and promising subset for testing (35). deepNGS Navigator can help in this selection process by projecting the sequences onto 2D maps, providing insights into their groups and diversity. We analyzed a synthetic, ML generated sequence set organized in a hierarchical tree structure rooted on a ”seed” sequence (24). Our method produced an embedding map that separated sequences by their origin generative method and captured the edit distance relationships more accurately than other approaches. We suggest embedding ML-generated sequences alongside repertoire data to guide selection, with a focus on overlapping clusters and their trajectories to identify natural and promising candidates.

deepNGS Navigator only needs sequence data as input, without needing prior information on V(D)J gene annotation. This is very useful in species like camelids where germline database is incomplete (2), or rabbits that affinity-matured sequences follow complex gene conversion paths (3). Additionally, unlike general repertoire mining methods that explore a relatively limited space of NGS sequences (36), deepNGS Navigator provides a broad overview of a large set of sequences, making it a flexible, easy-to-use tool for a wide range of contexts. Unlike clonotyping and phylogenetic trees, which rely heavily on accurate germline information and can produce unreliable results if the data is incorrect (5; 6), deepNGS Navigator using language models derives insights directly from sequence which makes it more robust to such challenges. For example, we used it to cluster naive and memory B-cell populations (25) and demonstrated clear discrimination between sequences based on their B-cell types. Additionally, the final map showed that sequences positioned close to each other in the embedding map are often biologically related based on their gene assignment, further validating the tool’s ability to capture meaningful relationships. The ability to analyze large-scale datasets and independence from gene annotation, makes deepNGS Navigator a practical solution for inferring clonal families as well. In this context, the input to deepNGS Navigator consists of nucleotides, resulting in sequences that are three times longer but reducing positional diversity from 20 possible amino acids to just 4 nucleotides. We benchmarked deepNGS using an in-silico dataset specifically designed for evaluation purposes and found that it accurately identified family lineages similar to state-of-the-art tools. Moreover, deepNGS outperformed existing methods in correctly separating noise from true lineage data. One application of deepNGS embedding maps is similar to tree-based methods, for identifying evolutionary neighbors of specific clones. We demonstrated that the coordinates on the embedding map can be used both to directly query such neighbors or to construct phylogeny-like trees.

As a promising future direction, we believe that combining sequence-based methods like deepNGS Navigator with structural insights could address some of the limitations inherent in sequence-only approaches. Such hybrid approach could provide a deeper understanding of B-cell evolution and complement ongoing experimental efforts to understand the mechanisms of B-cell affinity maturation (36). While this work focuses on the sequence-based clustering, the integration of structural insights will be the focus of future work on deepNGS Navigator. Finally, we hope that scientists and researchers, will find this tool useful and contribute to its ongoing development. Enhancements such as improving the interactive panel platform by adding features like motif search or filtration could greatly enrich its capabilities.

## DATA AVAILABILITY

The deepNGS Navigator source code is publicly available at: github.com/prescient-design/deepngs-navigator and github.com/prescient-design/deepngs-navigator-panel-app.

## References

1. E. Gallo, “The rise of big data: deep sequencing-driven computational methods are transforming the landscape of synthetic antibody design,” Journal of Biomedical Science, vol. 31, mar 2024.

2. L. Hanke, D. J. Sheward, A. Pankow, L. P. Vidakovics, V. Karl, C. Kim, E. Urgard, N. L. Smith, J. Astorga-Wells, S. Ekström, J. M. Coquet, G. M. McInerney, and B. Murrell, “Multivariate mining of an alpaca immune repertoire identifies potent cross-neutralizing sars-cov-2 nanobodies,” Science Advances, vol. 8, mar 2022.

3. J. J. Lavinder, K. H. Hoi, S. T. Reddy, Y. Wine, and G. Georgiou, “Systematic characterization and comparative analysis of the rabbit immunoglobulin repertoire,” PLoS ONE, vol. 9, p. e101322, jun 2014.

4. N. Nouri and S. H. Kleinstein, “A spectral clustering-based method for identifying clones from high-throughput b cell repertoire sequencing data,” Bioinformatics, vol. 34, p. i341–i349, jun 2018.

5. A. Yermanos, V. Greiff, N. J. Krautler, U. Menzel, A. Dounas, E. Miho, A. Oxenius, T. Stadler, and S. T. Reddy, “Comparison of methods for phylogenetic b-cell lineage inference using time-resolved antibody repertoire simulations (absim),” Bioinformatics, vol. 33, p. 3938–3946, aug 2017.

6. A. D. Yermanos, A. K. Dounas, T. Stadler, A. Oxenius, and S. T. Reddy, “Tracing antibody repertoire evolution by systems phylogeny,” Frontiers in Immunology, vol. 9, oct 2018.

7. O. Lindenbaum, N. Nouri, Y. Kluger, and S. H. Kleinstein, “Alignment free identification of clones in b cell receptor repertoires,” Nucleic Acids Research, vol. 49, p. e21–e21, dec 2020.

8. L. McInnes, J. Healy, and J. Melville, “Umap: Uniform manifold approximation and projection for dimension reduction,” 2018.

9. J. Devlin, M.-W. Chang, K. Lee, and K. Toutanova, “Bert: Pre-training of deep bidirectional transformers for language understanding,” 2018.

10. T. Chen, S. Kornblith, M. Norouzi, and G. Hinton, “A simple framework for contrastive learning of visual representations,” 2020.

11. J. N. Böhm, P. Berens, and D. Kobak, “Unsupervised visualization of image datasets using contrastive learning,” 2022.

12. V. A. Traag, L. Waltman, and N. J. van Eck, “From louvain to leiden: guaranteeing well-connected communities,” Scientific Reports, vol. 9, mar 2019.

13. “Github - genentech/absolve: Absolve antibody variable domain sequence analysis.” https://github.com/Genentech/Absolve, 2024. Accessed: 2024-10-02.

14. F. Zong, C. Long, W. Hu, S. Chen, W. Dai, Z.-X. Xiao, and Y. Cao, “Abalign: a comprehensive multiple sequence alignment platform for b-cell receptor immune repertoires,” Nucleic Acids Research, vol. 51, p. W17–W24, may 2023.

15. A. Honegger and A. Plückthun, “Yet another numbering scheme for immunoglobulin variable domains: An automatic modeling and analysis tool,” Journal of Molecular Biology, vol. 309, p. 657–670, jun 2001.

16. H.-L. Li, Y.-H. Pang, and B. Liu, “Bioseq-blm: a platform for analyzing dna, rna and protein sequences based on biological language models,” Nucleic Acids Research, vol. 49, p. e129–e129, sep 2021.

17. T. H. Olsen, I. H. Moal, and C. M. Deane, “Ablang: an antibody language model for completing antibody sequences,” Bioinformatics Advances, vol. 2, jan 2022.

18. “Compute a and b via mindist · github.” https://gist.github.com/NikolayOskolkov/9d00868443423063f8d9036a31c04f37, 2024. Accessed: 2024-10-20.

19. “Github - scipy/scipy: Scipy library main repository.” https://github.com/scipy/scipy, 2024. Accessed: 2024-10-12.

20. “Github - awslabs/amazon-denseclus: Clustering for mixed-type data.” https://github.com/awslabs/amazon-denseclus, 2024. Accessed: 2024-10-02.

21. “Github - holoviz/holoviz: High-level tools to simplify visualization in python..” https://github.com/holoviz/holoviz, 2024. Accessed: 2024-10-02.

22. M. Minot and S. T. Reddy, “Meta learning addresses noisy and under-labeled data in machine learning-guided antibody engineering,” Cell Systems, jan 2024.

23. L. van der Maaten and G. Hinton, “Visualizing data using t-sne,” Journal of Machine Learning Research, vol. 9, no. 86, pp. 2579–2605.

24. L. Li, E. Gupta, J. Spaeth, L. Shing, R. Jaimes, E. Engelhart, R. Lopez, R. S. Caceres, T. Bepler, and M. E. Walsh, “Machine learning optimization of candidate antibody yields highly diverse sub-nanomolar affinity antibody libraries,” Nature Communications, vol. 14, jun 2023.

25. “Different b cell subpopulations show distinct patterns in their igh repertoire metrics — elife.” https://elifesciences.org/articles/73111, 2024. Accessed: 2024-10-03.

26. K. Wang, X. Hu, and J. Zhang, “Fast clonal family inference from large-scale b cell repertoire sequencing data,” Cell Reports Methods, vol. 3, p. 100601, oct 2023.

27. N. Abdollahi, L. Jeusset, A. L. De Septenville, H. Ripoche, F. Davi, and J. S. Bernardes, “A multi-objective based clustering for inferring bcr clonal lineages from high-throughput b cell repertoire data,” PLOS Computational Biology, vol. 18, p. e1010411, aug 2022.

28. D. K. Ralph and F. A. Matsen, “Likelihood-based inference of b cell clonal families,” PLOS Computational Biology, vol. 12, p. e1005086, oct 2016.

29. U. Hershberg and E. T. Luning Prak, “The analysis of clonal expansions in normal and autoimmune b cell repertoires,” Philosophical Transactions of the Royal Society B: Biological Sciences, vol. 370, p. 20140239, sep 2015.

30. J. L. Bentley, “Multidimensional binary search trees used for associative searching,” Communications of the ACM, vol. 18, p. 509–517, sep 1975.

31. R. A. Norman, F. Ambrosetti, A. M. J. J. Bonvin, L. J. Colwell, S. Kelm, S. Kumar, and K. Krawczyk, “Computational approaches to therapeutic antibody design: established methods and emerging trends,” Briefings in Bioinformatics, vol. 21, p. 1549–1567, oct 2019.

32. A. Mahendra, A. Haque, P. Prabakaran, B. C. Mackness, T. P. Fuller, et al., “Honing-in antigen-specific cells during antibody discovery: a user-friendly process to mine a deeper repertoire,” Communications Biology, vol. 5, oct 2022.

33. K. Fischer, A. Lulla, T. Y. So, P. Pereyra-Gerber, M. I. J. Raybould, T. N. Kohler, J. C. Yam-Puc, T. S. Kaminski, R. Hughes, G. L. Pyeatt, F. Leiss-Maier, P. Brear, N. J. Matheson, C. M. Deane, M. Hyvönen, J. E. D. Thaventhiran, and F. Hollfelder, “Rapid discovery of monoclonal antibodies by microfluidics-enabled facs of single pathogen-specific antibody-secreting cells,” Nature Biotechnology, aug 2024.

34. A. A. R. Teixeira, S. D’Angelo, M. F. Erasmus, C. Leal-Lopes, F. Ferrara, L. P. Spector, L. Naranjo, E. Molina, T. Max, DeAguero, K. Perea, S. Stewart, R. A. Buonpane, H. G. Nastri, and A. R. M. Bradbury, “Simultaneous affinity maturation and developability enhancement using natural liability-free cdrs,” mAbs, vol. 14, sep 2022.

35. J. Kim, M. McFee, Q. Fang, O. Abdin, and P. M. Kim, “Computational and artificial intelligence-based methods for antibody development,” Trends in Pharmacological Sciences, vol. 44, p. 175–189, mar 2023.

36. Y.-C. Hsiao, H. A. Wallweber, R. G. Alberstein, Z. Lin, C. Du, A. Etxeberria, T. Aung, Y. Shang, D. Seshasayee, F. Seeger, M. Watkins, D. V. Hansen, C. J. Bohlen, P. L. Hsu, and I. Hötzel, “Rapid affinity optimization of an anti-trem2 clinical lead antibody by cross-lineage immune repertoire mining,” Nature Communications, vol. 15, sep 2024.

